# A molecular dynamic investigation of Human Rhinovirus 3C Protease Drug Target: Insights towards the design of potential Inhibitors

**DOI:** 10.1101/2022.03.06.483152

**Authors:** Ndumiso M Buthelezi, Daniel G. Amoako, Anou M. Somboro, Rene B. Khan, Hezekiel M. Kumalo

**Affiliations:** Discipline of Medical Biochemistry, School of Laboratory Medicine and Medical Science, University of KwaZulu-Natal, Durban, South Africa; Biomedical Resource Unit, College of Health Sciences, University of KwaZulu-Natal, Durban, South Africa

**Keywords:** 3C Protease, Rupintrivir, SG85, Human Rhinovirus, Molecular Dynamics, MM/GBSA

## Abstract

The 3C protease is distinguished from most proteases due to the presence of cysteine nucleophile that plays an essential role in viral replication. This peculiar structure encompassed with its role in viral replication has promoted 3C protease as an interesting target for therapeutic agents in the treatment of diseases caused by human rhinovirus (HRV). Herein we present a comprehensive molecular dynamics study of the comparison of two potent inhibitors, sg85 and rupintrivir complexed with HRV-3C protease. The binding free energy studies revealed a higher binding affinity for sg85 −58.853 kcal/mol than for rupintrivir −54.0873 kcal/mol and this was found to be in correlation with the experimental data. The energy decomposition analysis showed that, residues Leu 127, Thr 142, Ser 144, Gly 145, Tyr 146, Cys 147, His 161, Val 162, Gly 163, Gly 164, Asn 165, Phe 170 largely contributed to the binding of sg85, whereas His 40, Leu 127 and Gly 163 impacted the binding of rupintrivir. It further showed that His 40, Glu 71, Leu 127, Cys 147 Gly 163 and Gyl 164 are crucial residues that play a key role in ligand-enzyme binding; amongst these residues are residues of the conserved active site (His 40, Glu 71 and Cys 147). These findings provide a comprehensive understanding of the dynamics and structural features and will serve as guidance in the design and development of potent novel inhibitors of HRV.

**Graphical Abstract:** 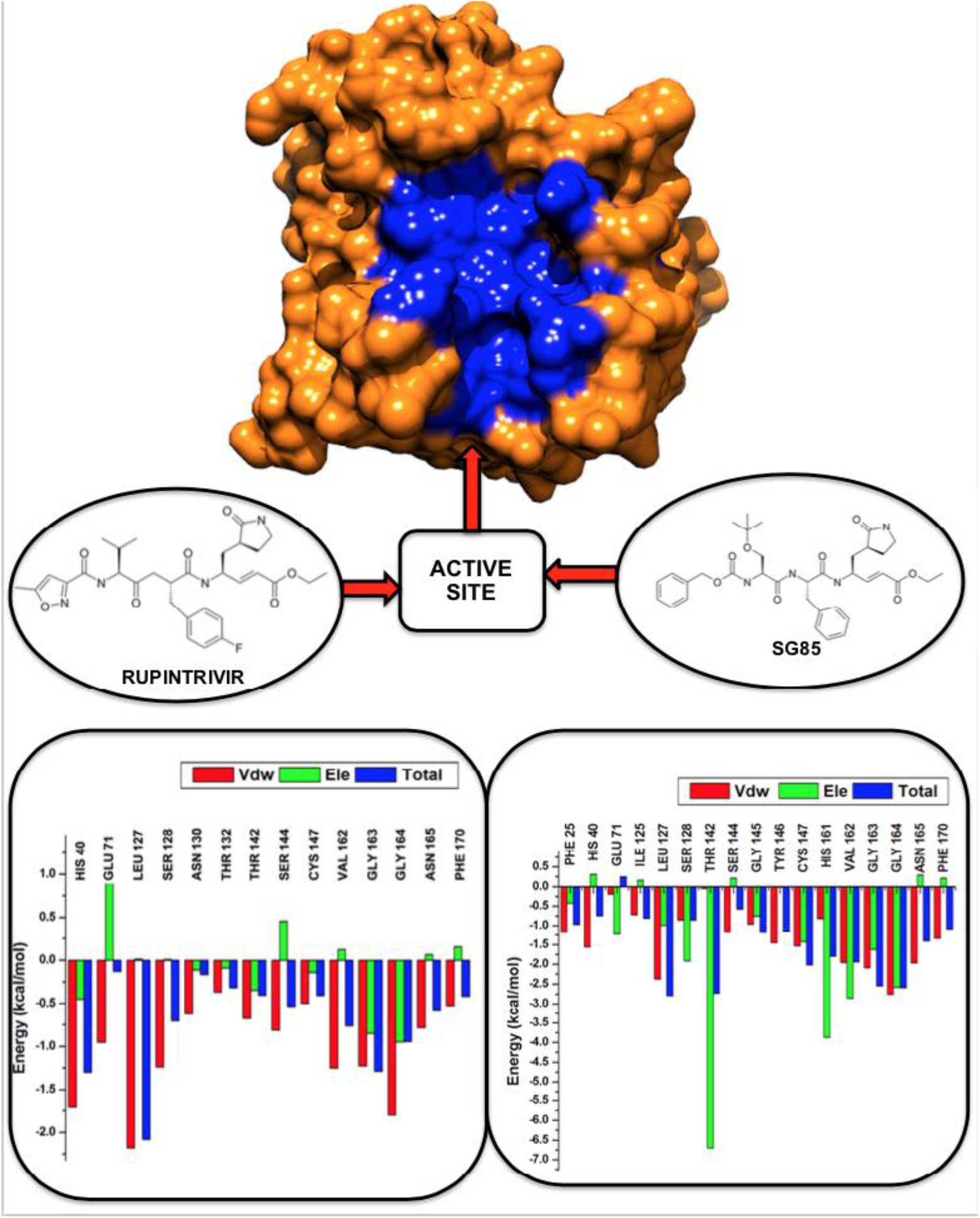

## 1. Introduction

Human rhinovirus (HRV), also known as the causative agent of common cold [1,2], are non-enveloped viruses that contain a single-strand ribonucleic acid (ssRNA) genome enclosed in an icosahedral (20-sided) capsid [3]. It is estimated that 18.8 billion and 150 million people were infected with upper respiratory and lower respiratory infections, respectively [4]. HRV is also reported by the World Health Organization (WHO) to be responsible for approximately 2 million children deaths necessitating the search and improvement of new potent drugs.

HRV is categorized into 3 species, namely, RV-A, RV-B, and RV-C [5]. RV-C being recently identified and studies show that HRV-3C-is most prevalent HRV identified in hospitalized children [6]. HRV is a positive-sense ssRNA virus of approximately 7200 bp [7]. Upon infection, the positive-strand RNA genome of HRV-3C is translated into a large poly-protein that is required for the production of new infectious virion and is dependent on two virally encoded proteases 2A and 3C. [8,9]. The 3C protease plays an indispensable role in viral replication in proteolytic cleavage of large polyproteins to functional proteins, and enzymes required for viral RNA replication both structurally and enzymatically [10]. The enzyme possess a strictly conserved catalytic active site as shown in **Fig.1 in blue**, thus, making it an attractive target for therapeutic agents in the treatment of diseases caused by human rhinovirus [11,12].

**Fig.1.**
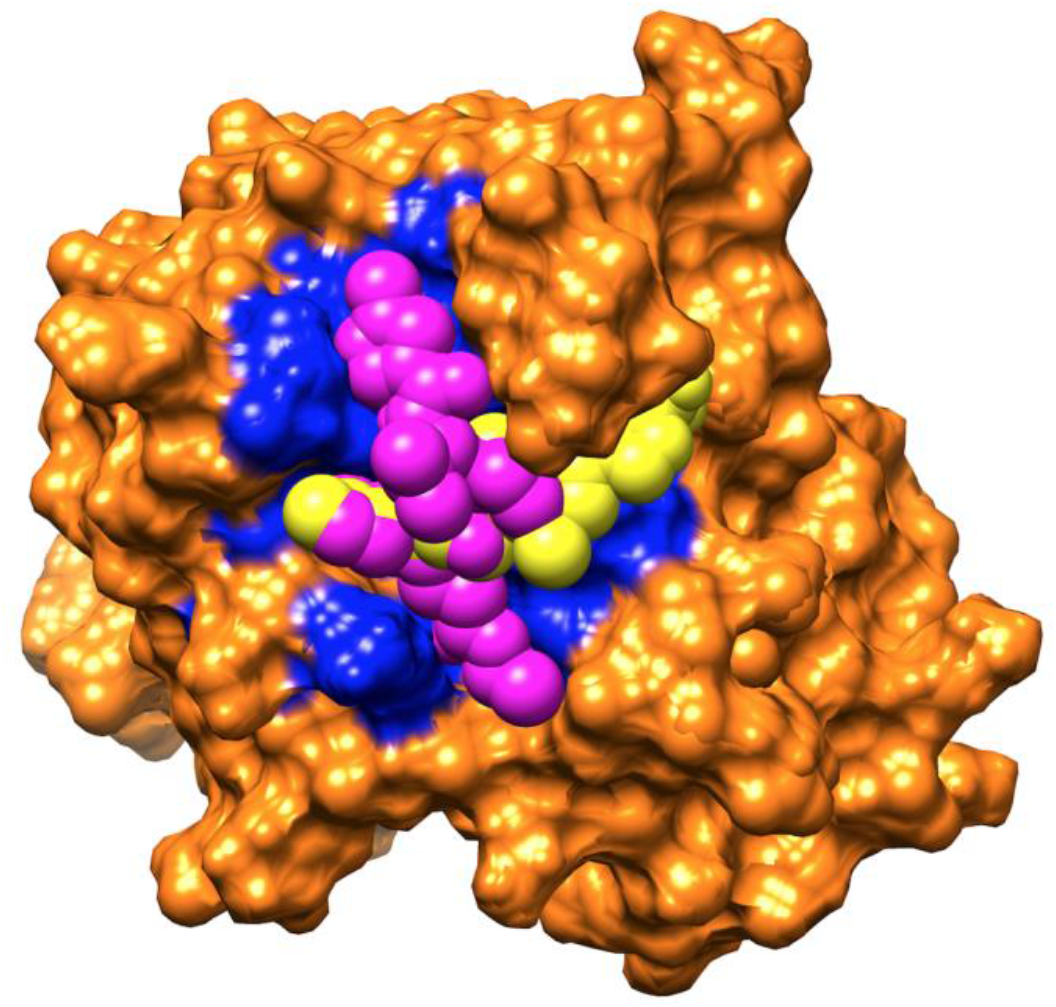
Depiction of Rupintrivir (Magenta) and Sg85 (Yellow) in the conserved active site (Blue). Several small molecule HRV inhibitors have been identified, with 3C protease (3CP) emerging as a promising therapeutic target [12]. It has recently been established that sg85 and rupintrivir inhibits HRV-3C protease.

Rupintrivir (formerly AG7088) is an irreversible inhibitor of the 3C protease of the human rhinovirus (HRV) that was discovered using structure-based drug design techniques [13]. Clinical trials reports showed that therapeutic application of rupintrivir was able to reduce viral load and moderate the severity of illness in a human experimental HRV challenge trial, hence providing a proof of concept for the mechanism of 3C protease inhibition. It was later discovered that rupintrivir had no significant effect on virus replication or disease severity, as such rupintrivir’s clinical development was halted. [14].

Sg85 is a peptidomimetics with Michael acceptor warheads permanently disable the protease by covalent binding to its catalytic site and is the most promising candidate [15]. In cell-based tests, Sg85 suppresses the replication of enteroviruses by inhibiting the EV68 3C-protease. [16].

It is believed that inhibition of picornavirus replication reduces the severity and shortens the duration of cold symptoms while rupintrivir is an irreversible inhibitor of HRV-3C protease and is regarded as the most potent novel peptidomimetic with inhibition activity against HRVs [17]. However, in terms of binding efficiency at the molecular level, there is no proof or comparison between these two inhibitors. Hence, the molecular understanding of HRV-3C protease structure and ligand-binding mechanisms remains unclear. Molecular dynamics simulations are one of the useful tools that have the capacity to provide a concise understanding at atomic level of complex systems [18]. To study the binding of the two inhibitors with HRV-3C protease, we employed molecular dynamics simulations (MD). In recent years, different post-dynamics analyses such as root mean square fluctuation, principal component analysis, Mechanics/Generalized Born Surface Area and dynamic cross-correlation analyses have been applied extensively in understanding the dynamics of biological complex systems at the atomic level. Therefore, this work provides understanding of molecular dynamics of HRV-3C protease complexed with sg85 and rupintrivir from a computational point of view. Furthermore, we aim at providing insight into the binding-landscape of the two inhibitors in HRV-3C protease by conducting a broad comparative MD analysis of the HRV-3C-Free, HRV-3C-Sg85 and HRV-3C-Rupintrivir systems. We hope that this work could serve as a cornerstone in drug design and developments of more potential drugs against HRV protease, which could lead the fight against the virus.

## 2. Computational Methodology

### 2.1. System Preparations

The X-ray crystal structure of human rhinovirus-3C protease, PDB code **5FX6**, bound to rupintrivir was extracted from Protein Bank Database (www.rcsb.org) [19].The Michael acceptor, Sg85, was obtained from the PDB 2YNB where it was in complex with corona virus HKU4[20]. Prior to molecular docking all non-standard residues including ions, Na+, cl-, H_2_O were removed from the structure of human rhinovirus-3C protease. Subsequently, Sg85 was removed from the 2YNB complex and optimized using Avogadro software before docking into the binding site of human rhinovirus 3C protease using Autodock vina integrated in UCSF Chimera. All missing residues were remodeled using the modeler plugin incorporated in UCSF Chimera graphical user interface [21]. In preparation for MD simulation, hydrogen atoms and amber charges were then added to the ligand and saved separately in mol2 format whiles the protein was saved in pdb format. Molecular visualization of the Ligand and receptor were conducted in Chimera and Avogadro softwares [22].

### 2.2. Molecular Dynamic Simulation

The simulation setup was done using the Graphical Processing Unit (GPU) version of Particle Mesh Ewald Molecular Dynamics (PMEMD) integrated in the Amber 14 suite [23,24]. Atomic partial charges were generated for the ligands using the General Amber Force Field GAFF force field during the ANTECHAMBER stage on the AMBER 14 [25]. Likewise, the FF14SB force field was used to parametize the receptor [25] [26]. This was then followed by the LEAP phase, where additional hydrogen atoms were added to the system. Also, neutralization of the system was carried out through the addition of counter ions such as NA^+^ and Cl^-^. The system was then subjected to solvation by immersing in an orthorhombic box with TIP3P [27] water molecules at a distance of 10 Å. In our previous work, system setup, minimizations, heating and equilibration steps were thoroughly explained [28,29]. Subsequently the system was subjected to a 200ns MD simulation time, and all trajectories generated during the simulation run were then saved for every 1ps and analyzed. The CPPTRAJ and PTRAJ modules [30] incorporated in the Amber 14 suite were used for analyzing the trajectories: Parameters such as RMSD, RMSF, RoG, DCC, PCA were determined. The Origin data analysis tool (www.originlab.com) was utilized for all graphical plots while Chimera [21] was used for visualizations.

### 2.3. Binding energy calculations

Molecular Mechanics/Generalized Born Surface Area (MM/GBSA) was used to calculate the binding free-energy profiles of rupintrivir and sg85 bound to HRV-3C protease. [31,32]. The binding free energy calculation gives insights into the association of the protein-ligand in a complex and also gives an account on the end point energy calculation. Diffirential binding free energy was calculated taking into account 1000 snapshots from 200 ns trajectories. The calculation of the binding free energy is described by the following set of equations:

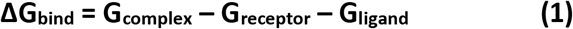

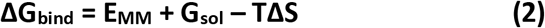

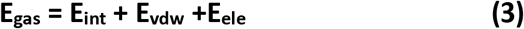

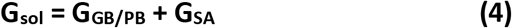

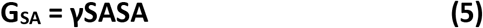

Where E_gas_ represents gas-phase energy, internal energy is represented by E_int_, whiles E_ele_ and E_vdw_ represent the electrostatic and Van der Waals contributions, respectively. E_gas_ is the gas phase, which is directly elevated from the FF14SB force terms. G_sol_, which stands for solvation energy, can be divided into polar and nonpolar contribution states. The polar solvation contribution, G_GB_, is calculated by solving the GB equation, whereas the nonpolar solvation contribution, a water probe radius of 1.4 Å is used to determine G_SA_, is estimated using a. T and S represent temperature and total solute entropy, respectively. A per-residue decomposition analysis of the interaction energy for each residue was performed using the MM/GBSA method to determine the contribution of each amino acid to the total binding free energy profile between rupintrivir and sg85.

### 2.4. Dynamic Cross-correlation Analysis (DCC)

The dynamic cross correlation (DCC) method has been widely used to calculate the correlation coefficients of motions between atoms in a protein. [33]. The CPPTRAJ module was used to calculate the residue-based fluctuations during the simulation [30]. DCC is represented by the following formula given below:

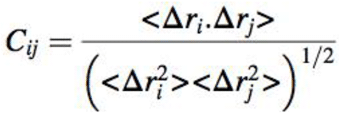

With C_ij_ being the cross-correlation coefficient, which varies with a range of −1 to +1. A fully correlated and anti-correlated motion represents the upper and lower limits during the simulation, respectively. Where i and j represent the i^th^ and j^th^ residues and Δr_i_ and Δ_i_ symbolize displacement vectors correspond to ith and jth residues, respectively. In this study, DCC calculations were carried out considering the backbone C-α atomic fluctuations.

### 2.4. Principal Component Analysis (PCA)

Principal component analysis (PCA) also referred to as essential dynamics (ED) and is one of the utmost unconventional methods for trajectory analysis. PCA has proven to be powerful and robust, opening up new avenues for visualizing and exploring the dynamics of protein cavities. [34,35]. PCA describes the eigenvectors and eigenvalues, which represent the direction of motions and the amplitudes in those directions of the protein, respectively [36]. Following the removal of the ions and solvent from the 200 ns MD trajectories. The first two principal components (PCA1 and PC2) were computed using the CPPTRAJ module in AMBER 14 on C-α atoms across 1000 snapshots at a time period of 100 ps. The Microcal Origin software was used to generate corresponding PCA scatter plots. (www.originlab.com).

## 3. Results and discussion

### 3.1. Root of mean square deviation (RMSD)

The root-mean-square deviation (RMSD) was plotted to identify the stability of studied enzyme-ligand complexes during the 200 ns MD trajectories. Fig.S1 highlights the RMSD of HRV-3C-Free, HRV-3C-Sg85 and HRV-3C-Rupintrivir.

Throughout the simulation, all three systems showed to have converged steadily and their stability can be described as acceptable. In Fig.S1, HRV-3C-Free maintained the highest average of 1.7643±0.03972 Å compared to HRV-3C-Rupintivir and HRV-3C-Sg85 systems with an average RMSD of 1.2541±0.1399 Å and 0.9714±0.0826 Å, respectively. A drastic fluctuation at 150 ns is of significant note, though the system soon regains stability at 170 ns. However, following the analysis conducted, a stabilization and convergence of the systems, all the systems had fluctuations of less than <2.5 Å.

### 3.2. Root Mean Square Fluctuation (RMSF)

Proteins are bound to undergo structural fluctuations when subjected to physiological conditions. Relatively high and low structural fluctuations are known to have a direct impact on protein function, and occur at a range of time scales during the simulation, in our case, nanoseconds [37]. A RMSF plot in Fig.S2 was used to analyze the amino acid fluctuations for all the HRV systems.

In the current work, a similar trend in fluctuations was noticed in HRV-3C-Free, HRV-3C-Rupitrivir and HRV-3C-Sg85 with an average RMSF values of 0.7518±0.3246 Å, 0.7144±0.3328 Å and 0.6098±0.1981 Å, respectively. Hence, HRV-3C-Free system displayed more flexibility compared to HRV-3C-Rupintrivir and HRV-3C-Sg85. The most significant change was observed in the active site residues in the regions 120-138 and 140-150 of the HRV-3C-Free enzyme and HRV-3C-Rupintrivir displayed higher fluctuations with reference to the HRV-3C-Sg85. In addition, the proximal residues in the region 25-35 of the HRV-3C-Free exhibit higher fluctuations compared to both HRV-3C-Rupintrivir and HRV-3C-Sg85. As evident in Fig.S2, Sg85 has reformed the overall protein flexibility compared to Rupintrivir.

Furthermore, Fig.S2 gives information about the β-hairpin in the active site residues in the regions 120-138 of all systems. It has been proposed that the β-hairpin acts as launch sites in initial events of protein folding [38]. Herein, the evolutions of the β-hairpin during a MD simulation for all conformations were examined and snapshots along the trajectories of MD simulations are shown in Fig.2.

**Fig.2.**
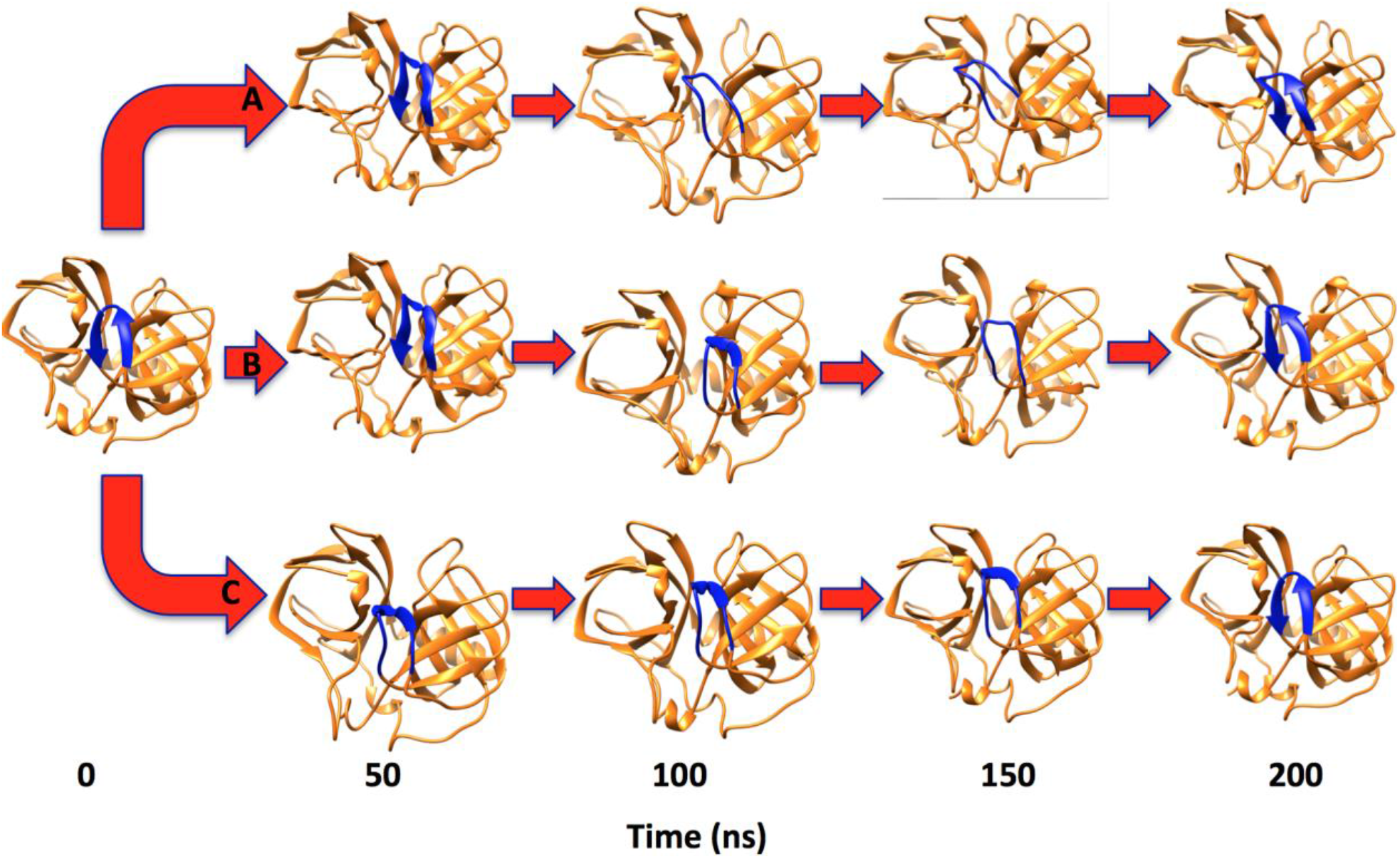
Snapshots of HRV-3C-Free (**A**), HRV-3C-Rupintrivir (**B**) and HRV-3C-Sg85 (**C**) conformations captured at 50 ns intervals throughout a 200 ns MD simulation.

During the simulation of the conformations, the β-hairpin on the HRV-3C-Free system disappeared at 100 ns and 150 ns with an increase in loop length, whereas the β-hairpin on the HRV-3C-Rupintrivir conformation disappeared at 150 ns with an increase *in loop length* (see Fig.S2). On the other hand, snapshots of the HRV-3C-Sg85 suggested a β-hairpin to be stable throughout a 200 ns MD simulation. Hence, these findings are correlated to those of RMSF presented herein, which justify a greater existence of fluctuations of HRV-3C-Free and HRV-3C-Rupintrivir compared to HRV-3C-Sg85 in the regions 120-138. Recent works suggested that the β-hairpin could form nucleation sites for protein folding, however, the molecular mechanisms by which proteins fold have not been yet fully understood [39–41]. As a result, the findings presented here could serve as basis for understanding the sequence and structural context of HRV-hairpin. Experimental results have shown Sg85 to be active against a wide range of HRV isolates suggesting that it has a higher binding affinity for the binding site compared to Rupintrivir[42]. We can conclude from the results that location of Sg85 in the active site leads to a conformational rigidity and β-hairpin stability [15].

### 3.3. Radius of Gyration (Rg)

Radius of gyration (Rg) was computed to measure the compactness of the protein structure and additionally to give details into stability of the complexed systems [43,44]. An Rg plot in Fig.S3 was used to analyze the overall protein dimensions of the HRV systems.

From Fig.S3 the similarities between the three systems are obvious, as evidenced by the similarity in residual alignment within the secondary and tertiary structures. HRV-3C-Free and HRV-3C-Rupintrivir displayed a slighter higher Rg compared to HRV-3C-Sg85. These results are in accordance with RMSF, which justified a substantial increase in biomolecular flexibility of the HRV-Free and HRV-3C-Rupintrivir when compared to the HRV-3C-Sg85. This indicates that HRV possesses a rigid structural stability when it is bound to Sg85 as compared to. A theoretical explanation for this observation is that binding of Sg85 to the active site of HRV causes conformational rigidity, which inhibits HRV’s conformational flexibility.

### 3.4. Dynamic Cross Correlation Analysis (DCC)

The dynamic cross correlation was applied to quantify the correlation coefficients of motions of the C-α atom fluctuations between atoms of a protein. The different correlation motion of the systems was analyzed. DCC plots presented in Fig.3 are interpreted based on the different colors with yellow to red indicating a strong correlation while blue to black indicate a strong anti-correlation of specific residue movements.

**Fig.3:**
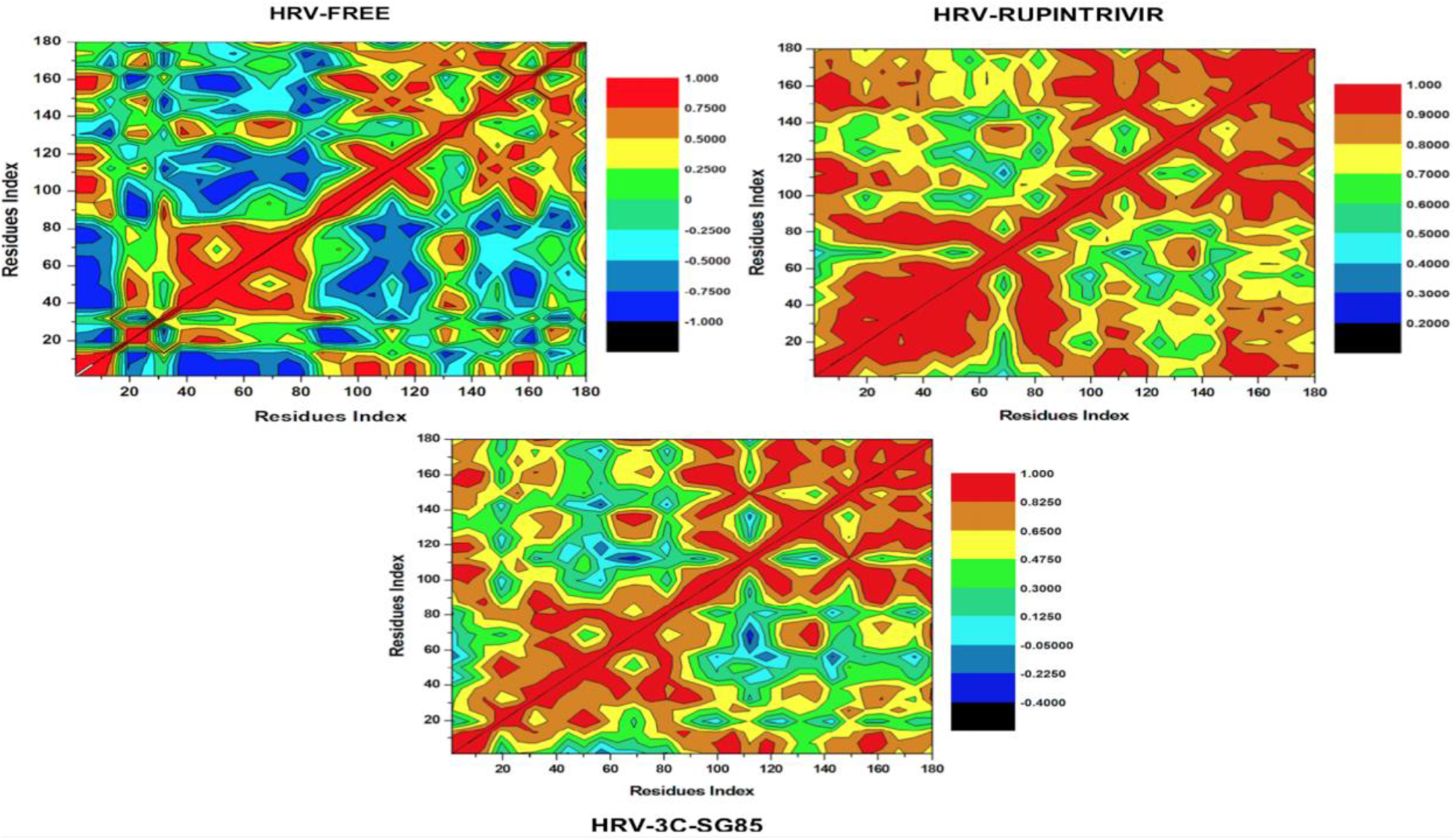
The dynamic cross-correlation matrix analyses during the 200 ns simulation for the HRV-3C-Free, HRV-3C-Rupintrivir and HRV-3C-Sg85

Anti-correlated residual motion in Fig.3, for and sg85 occur between residues 50 – 70 relative to 110, while a strong correlated residual motion in case of rupintrivir occur between 20 – 80 relative to each other. The correlation map displays a strong correlation of residual motion in the HRV-3C-Rupintrivir conformation, while in the HRV-3C-Sg85 conformation there was an observed greater anti-correlated motions (negative correlations) during simulation time. This is supported by the RMSF results, which also shows a strong interaction with residues compared to HRV-3C-Rupintrivir.

### 3.5. Principal Component Analysis (PCA)

Fig.4 highlights the most significant alterations in atomic motion across principal components of the HRV-3C-Free, HRV-3C-Rupintrivir and HRV-3C-Sg85. The conformational variations of sg85 and rupintrivir were characterized using PCA. PCA is a tool commonly used for conformational variations. Graphical plots of conformational motions of all systems were plotted on the two principal components (PC1 vs PC2), to provide a more detailed understanding of the conformational changes in the systems.

**Fig.4:**
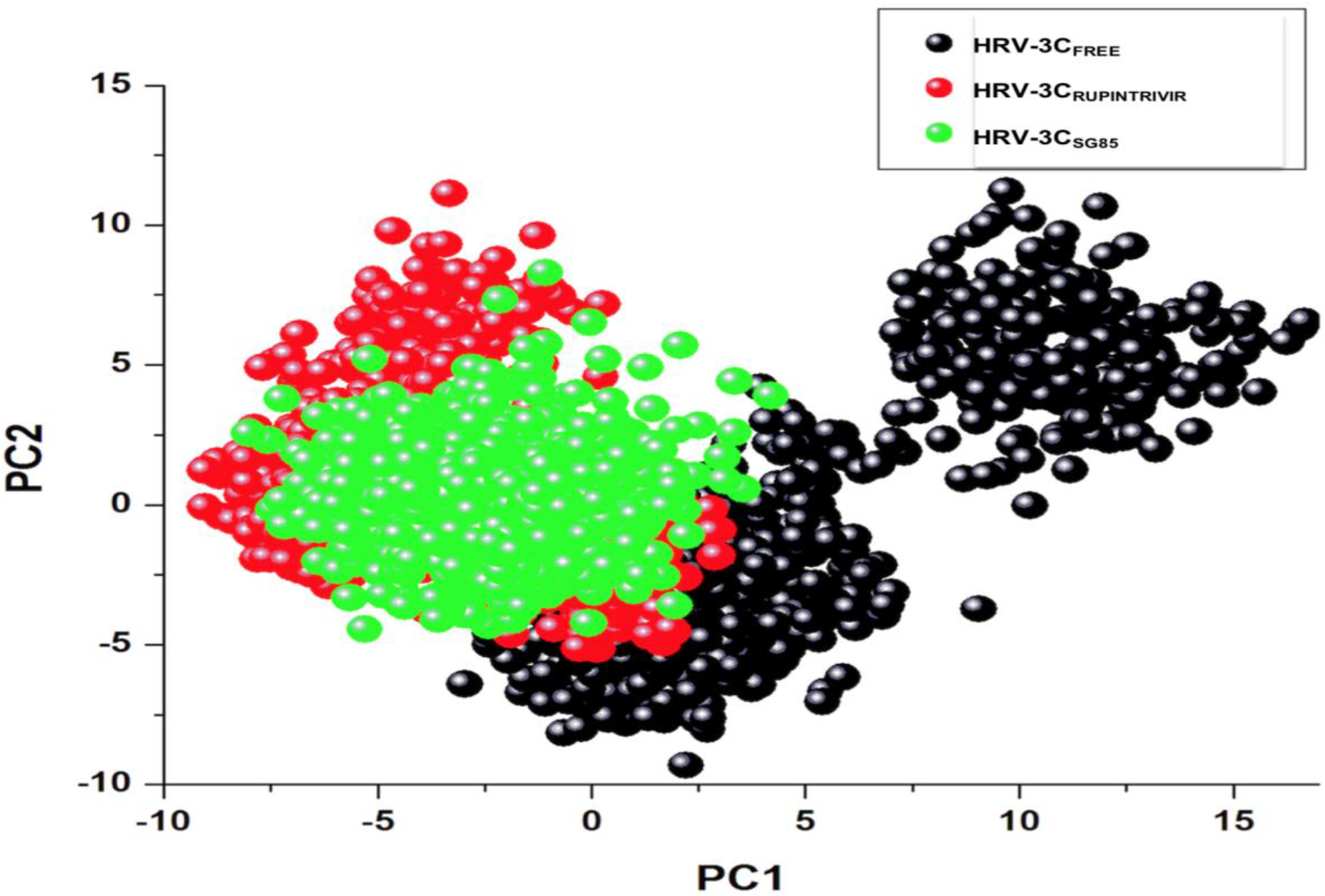
Projection of PC1 over PC2 for HRV-3C-Free, HRV-3C-Rupintrivir and HRV-3C-Sg85 conformations of HRV-3C.

Fig.4 illustrates that the Sg85 (in black) showed to have the largest surface area followed by Rupintrivir (in red) when compared to Free (in green). This could imply that the Sg85 is less flexible allowing a more easy binding toward the HRV enzyme compared to Rupintrivir. As evident in Fig.S2 and Fig.2, the HRV-3C-Free showed to be more flexible in contrast to Rupintrivir and Sg85. The evidence in Fig.4 unveils basic insights of the dynamic behavior of biological systems at molecular level.

### 3.6. MMPB-SA binding free energy calculation

The overall binding free energy for both Sg85 and Rupintrivir was calculated using the MM/PBSA method. Per-residue energy decomposition was computed to gain insight to the ligand-residue interaction and this was accomplished by 1000 snapshots of the 200 ns simulation trajectories.

As evident in table 1, the respective binding free energies (ΔG_binding_) for Sg85 and Rupintrivir was estimated to be **-58.853** kcal/mol and **-54.087** kcal/mol. The binding free energy of Sg85 is higher by (~4 kcal/mol) than that of Rupintrivir, which further suggests that Sg85 has a more intimate interaction with the active site residues compared to sg85. This is in great accordance with recent publications that have rated Sg85 as the most potent inhibitor of HRV enzyme [15].

**Table 1:**
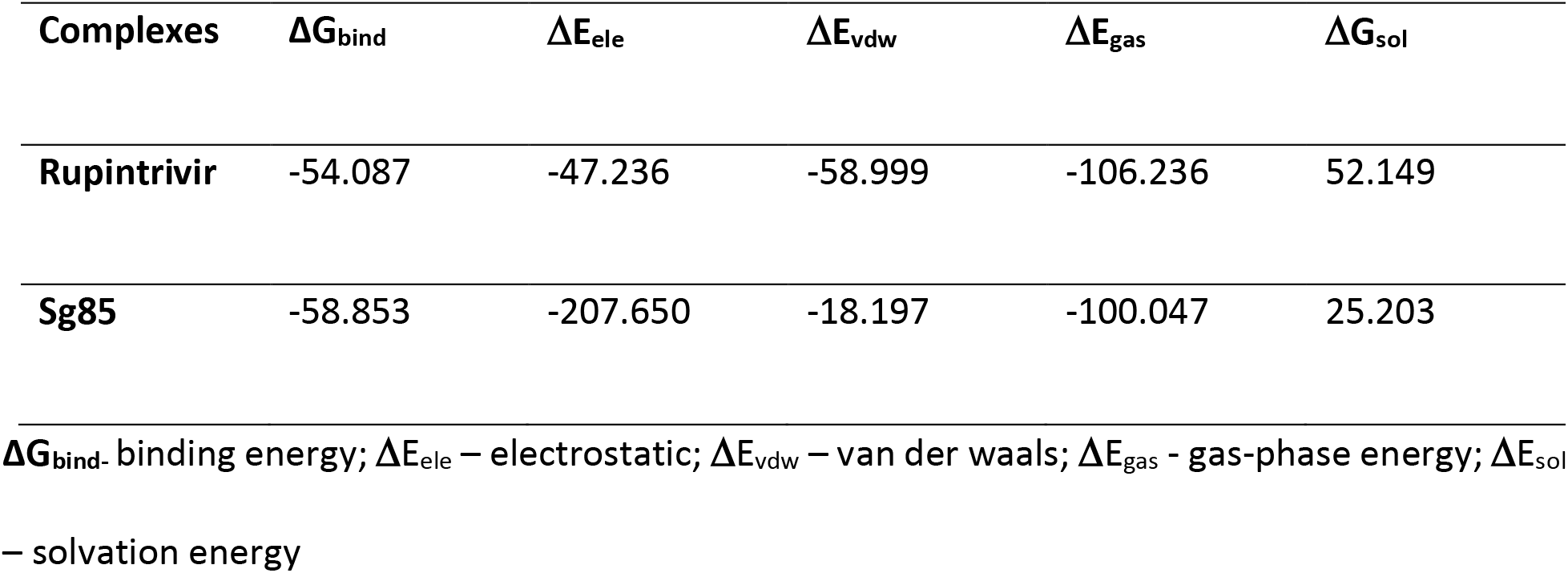
MM/GBSA based on binding free energy profile of Rupintrivir and Sg85 bound with 3C.

| Complexes | $\Delta G_{\text{bind}}$ | $\Delta E_{\text{ele}}$ | $\Delta E_{\text{vdw}}$ | $\Delta E_{\text{gas}}$ | $\Delta G_{\text{sol}}$ |
| --- | --- | --- | --- | --- | --- |
| Rupintrivir | -54.087 | -47.236 | -58.999 | -106.236 | 52.149 |
| Sg85 | -58.853 | -207.650 | -18.197 | -100.047 | 25.203 |

$\Delta G_{\text{bind}}$ . binding energy; $\Delta E_{\text{ele}}$ – electrostatic; $\Delta E_{\text{vdw}}$ – van der waals; $\Delta E_{\text{gas}}$ - gas-phase energy; $\Delta E_{\text{sol}}$ – solvation energy

As revealed from the per-residue energy contribution plots shown in Fig.5, The larger residual energy contributions (|ΔG_binding_ | > - 1) were **Leu 127, Thr 142, Ser 144, Gly 145, Tyr 146, Cys 147, His 161, Val 162, Gly 163, Gly 164, Asn 165, Phe 170 and His 40, Leu 127 and Gly 163,** for Rupintrivir and Sg85, respectively. However, in the case of sg85, **Glu 71**, a catalytic triad residue, showed an unfavorable energy contribution (+0.256 kcal/mol) and interestingly sg85 emerged as the potent inhibitor between the two. Moreover, both inhibitors interact with the key residues **His 40**, **Glu 71** and **Cys 147**. Thus, these results suggest that QSAR model should include these key residues for an efficient and effective drug design.

**Fig.5:**
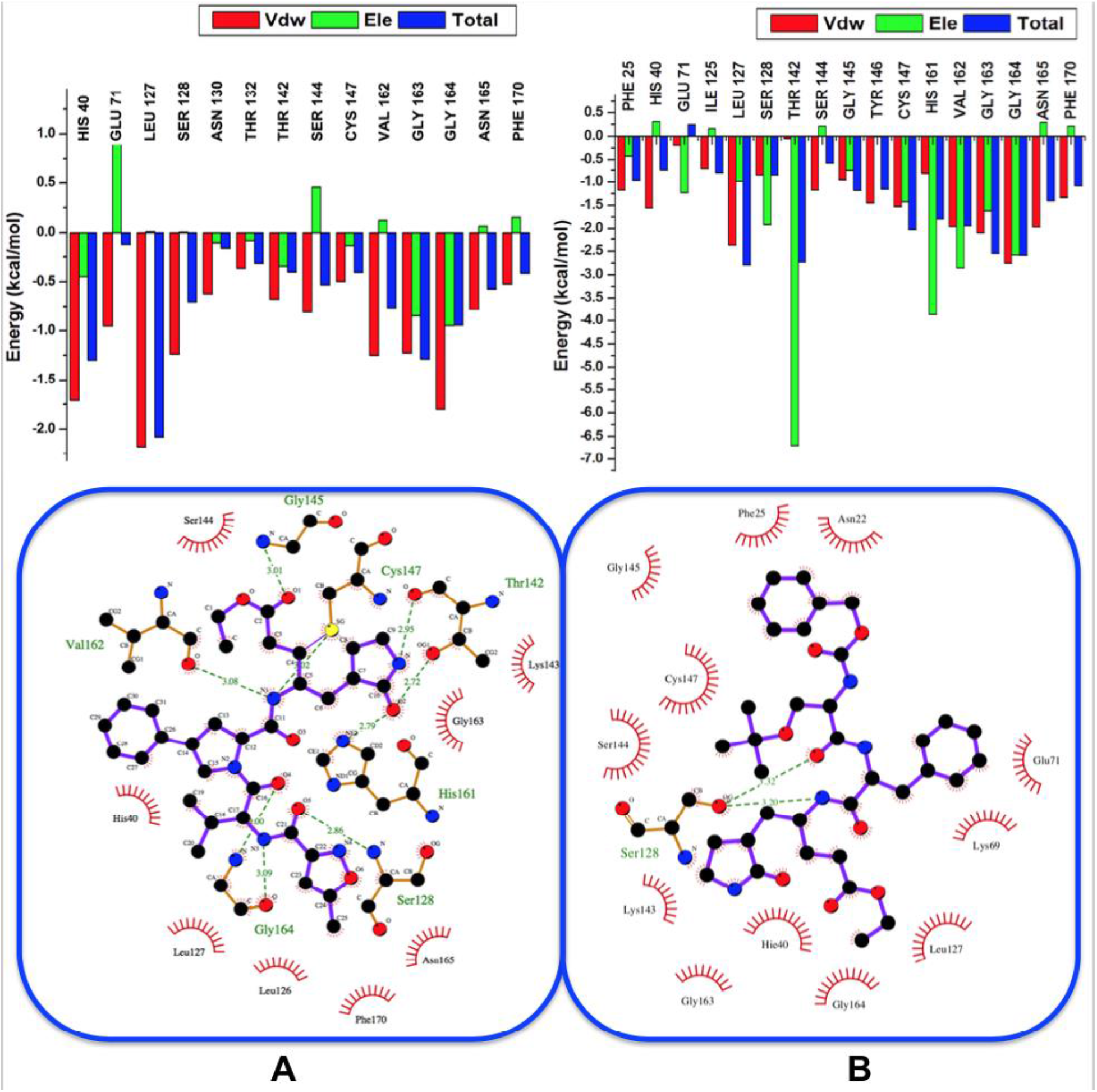
A diagrammatic representation of the per-residue graphs and 2D ligand interaction plots showing BFE contribution for (**A**) Rupintrivir and (**B**) Sg85 bound to HRV-3C.

The current study highlights the most important amino acids involved in inhibitor binding. The generated pharmacophore library contains only highly contributing amino acid residues, as determined by free binding energy contributions obtained from molecular dynamic (MD) simulations. [45,46]. We trust that the findings from this study serve as a foundation towards the design of new inhibitors that will interact with catalytic triad residues to discover novel compounds to inhibit HRV using a pharmacophore model.

## 4. Conclusion

Ligand-protein interaction and binding at molecular level is an imperative biological process. In this report, we have provided insights into the binding mechanism of the rupintrivir and sg85 on 3C protease with the application of advance molecular dynamics simulation tools. It has been experimentally reported that sg85 is a potent inhibitor when compared to rupintrivir but to this end, hence this comparative study was conducted using various advanced MD simulation and post-analysis tools to gain understanding into the binding landscape and dynamic structural features associated with the binding of rupintrivir and sg85 on 3C protease. From the results obtained, sg85 induced a more stable protein structure thus suggesting a more intimate binding and interaction of sg85 with the 3C protease when compared to rupintrivir as seen from the RMSD, RMSF, RG, DCC and PCA plots. The binding free energy calculation revealed a higher binding affinity for sg85 −58.853 kcal/mol than for rupintrivir −54.087 kcal/mol and this is in correlation with the experimental data. This thus suggests that sg85 can be used as scaffold to design new inhibitors. The energy decomposition analysis showed that, residues Leu 127, Thr 142, Ser 144, Gly 145, Tyr 146, Cys 147, His 161, Val 162, Gly 163, Gly 164, Asn 165, Phe 170 largely contributed to the binding of sg85, whereas His 40, Leu 127 and Gly 163 largely contributed to the binding of rupintrivir. The energy decomposition revealed that His 40, Glu 71, Leu 127, Cys 147 Gly 163 and Gyl 164 are crucial residues that play a key role in ligand-enzyme binding; amongst these residues are residues of the conserved active site (His 40, Glu 71 and Cys 147). Therefore, from these results we believe that the residue interactions and the binding free energy analysis must be considered for a pharmacophore model and structure-based design of novel and potent inhibitors of HRV.

## Acknowledgements

NMB acknowledges School of Health Sciences, UKZN, for financial assistance and the Center of High Performance Computing (CHPC), for computational facility (www.chpc.ac.za).

## Supplementary Material

**Figure S1:**
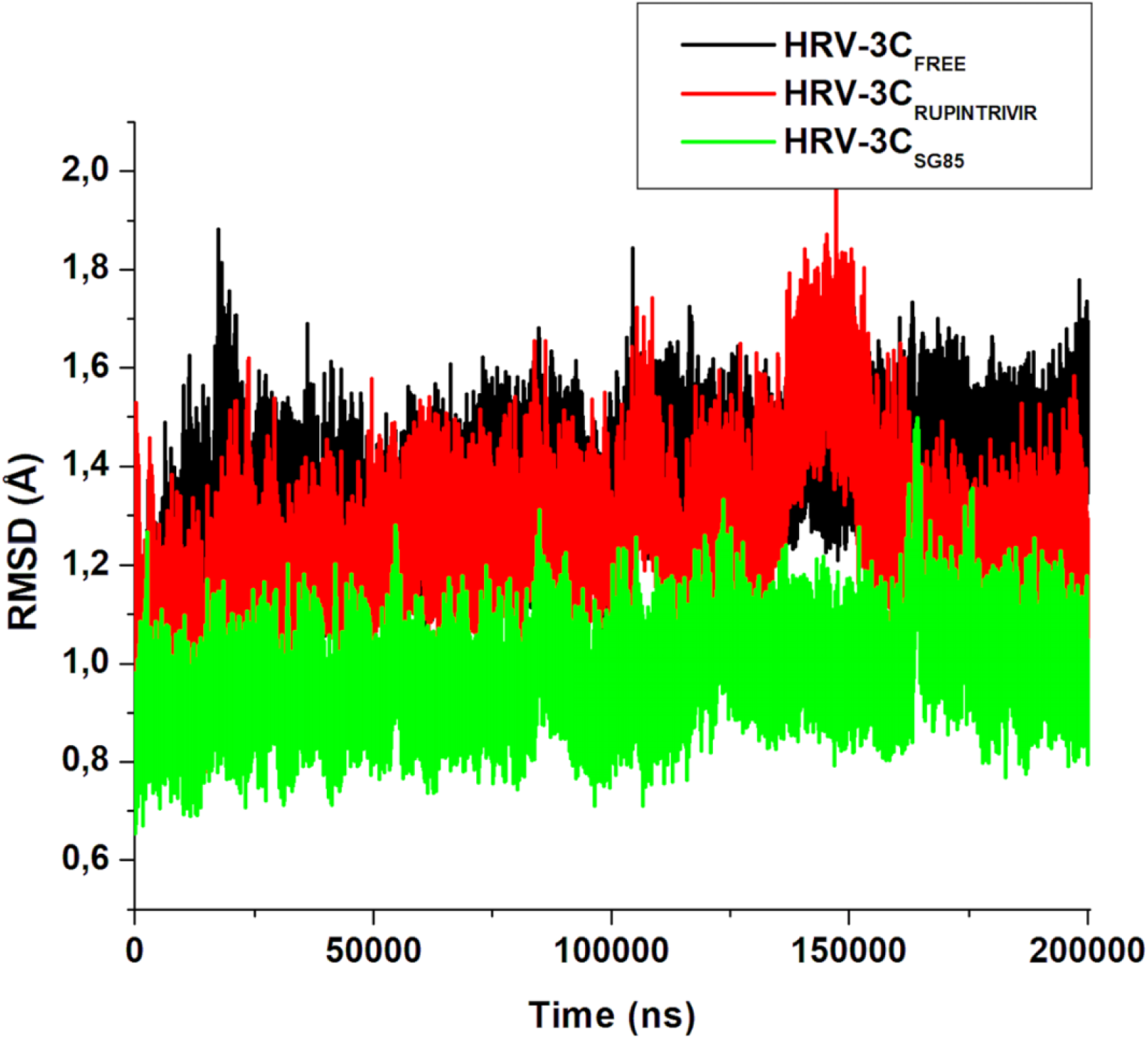
RMSD of minimized structures of HRV-3C-Free, HRV-3C-Rupintrivir and HRV-3C-Sg85 recorded over 200ns MD simulation.

**Figure S2:**
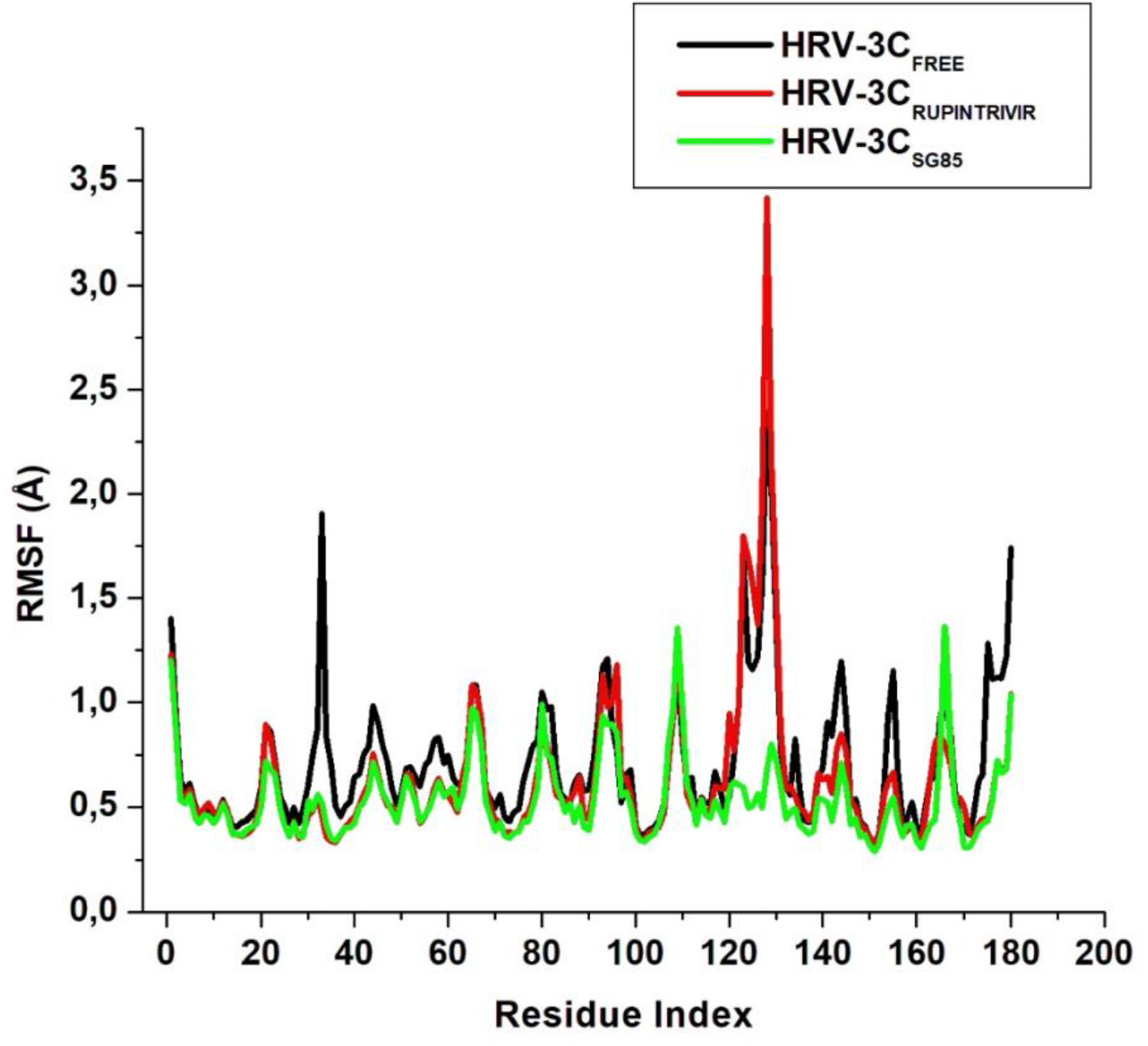
RMSF of minimized structures of HRV-3C-Free, HRV-3C-Rupintrivir and HRV-3C-Sg85 recorded over 200ns MD simulation.

**Figure S3:**
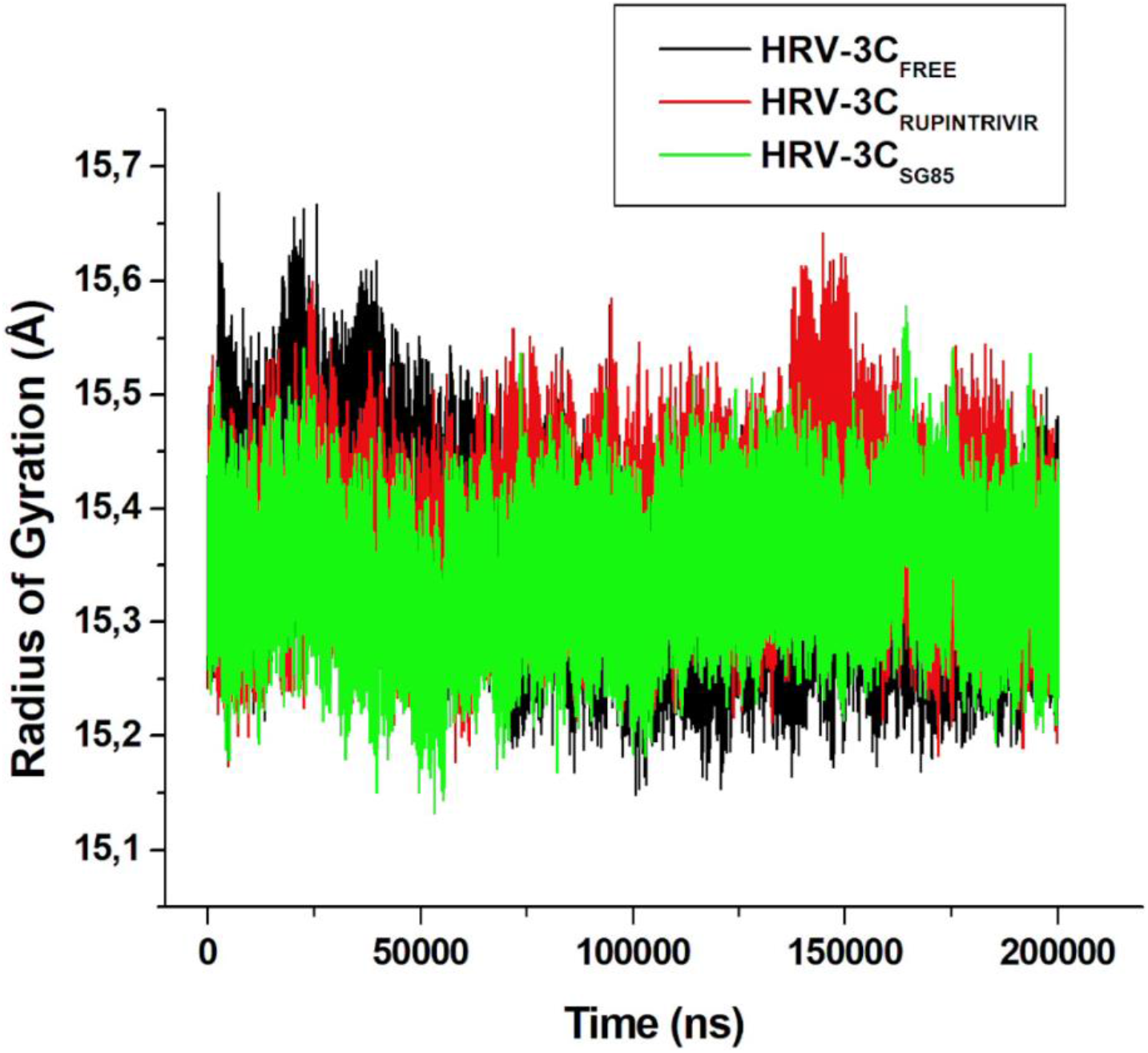
Rg of minimized structures of HRV-Free, HRV-3C-Rupintrivir and HRV-3C-Sg85 recorded over 200ns MD simulation.

## Reference

1. Atkinson, S.K.; Sadofsky, L.R.; Morice, A.H. How does rhinovirus cause the common cold cough? BMJ Open Respir. Res. 2016, 3.

2. Stobart, C.C.; Nosek, J.M.; Moore, M.L. Rhinovirus Biology, Antigenic Diversity, and Advancements in the Design of a Human Rhinovirus Vaccine. Front. Microbiol. 2017, 8, doi:10.3389/fmicb.2017.02412.

3. Palmenberg, A.C.; Gern, J.E. Classification and Evolution of Human Rhinoviruses. Methods Mol. Biol. 2015, 1221, 1–10, doi:10.1007/978-1-4939-1571-2_1.

4. Lamborn, I.T.; Jing, H.; Zhang, Y.; Drutman, S.B.; Abbott, J.K.; Munir, S.; Bade, S.; Murdock, H.M.; Santos, C.P.; Brock, L.G.; et al. Recurrent rhinovirus infections in a child with inherited MDA5 deficiency. J. Exp. Med. 2017, 214, 1949–1972, doi:10.1084/jem.20161759.

5. Kerr, S.-L.; Mathew, C.; Ghildyal, R. Rhinovirus and Cell Death. Viruses 2021, 13, 629, doi:10.3390/v13040629.

6. L’Huillier, A.; Kaiser, L.; Petty, T.; Kilowoko, M.; Kyungu, E.; Hongoa, P.; Vieille, G.; Turin, L.; Genton, B.; D’Acremont, V.; et al. Molecular Epidemiology of Human Rhinoviruses and Enteroviruses Highlights Their Diversity in Sub-Saharan Africa. Viruses 2015, 7, 6412–6423, doi:10.3390/v7122948.

7. Jacobs, S.E.; Lamson, D.M.; St. George, K.; Walsh, T.J. Human Rhinoviruses. Clin. Microbiol. Rev. 2013, 26, 135–162, doi:10.1128/CMR.00077-12.

8. Skorenski, M.; Sienczyk, M. Viral proteases as targets for drug design. Curr. Pharm. Des. 2013, 19, 1126–1153.

9. Flather, D.; Semler, B.L. Picornaviruses and nuclear functions: targeting a cellular compartment distinct from the replication site of a positive-strand RNA virus. Front. Microbiol. 2015, 6, doi:10.3389/fmicb.2015.00594.

10. Sun, D.; Chen, S.; Cheng, A.; Wang, M. Roles of the Picornaviral 3C Proteinase in the Viral Life Cycle and Host Cells. Viruses 2016, 8, 82, doi:10.3390/v8030082.

11. Jensen, L.M.; Walker, E.J.; Jans, D.A.; Ghildyal, R. Proteases of Human Rhinovirus: Role in Infection BT - Rhinoviruses: Methods and Protocols. In; Jans, D.A., Ghildyal, R., Eds.; Springer New York: New York, NY, 2015; pp. 129–141 ISBN 978-1-4939-1571-2.

12. Ullah, R.; Shah, M.A.; Tufail, S.; Ismat, F.; Imran, M.; Iqbal, M.; Mirza, O.; Rhaman, M. Activity of the Human Rhinovirus 3C Protease Studied in Various Buffers, Additives and Detergents Solutions for Recombinant Protein Production. PLoS One 2016, 11, e0153436, doi:10.1371/journal.pone.0153436.

13. Matthews, A.D.A.; Dragovich, P.S.; Webber, S.E.; Fuhrman, S.A.; Patick, A.K.; Hendrickson, T.F.; Love, R.A.; Prins, T.J.; Marakovits, J.T.; Zhou, R.; et al. Structure-Assisted Design of Mechanism-Based Irreversible Inhibitors of Human Rhinovirus 3C Protease with Potent Antiviral Activity against Multiple Rhinovirus Serotypes Source: Proceedings of the National Academy of Sciences of the United States of Amer. 2018.

14. Binford, S.L.; Weady, P.T.; Maldonado, F.; Brothers, M.A.; Matthews, D.A.; Patick, A.K. In Vitro Resistance Study of Rupintrivir, a Novel Inhibitor of Human Rhinovirus 3C Protease. Antimicrob. Agents Chemother. 2007, 51, 4366–4373, doi:10.1128/AAC.00905-07.

15. Lacroix, C.; George, S.; Leyssen, P.; Hilgenfeld, R.; Neyts, J. The Enterovirus 3C Protease Inhibitor SG85 Efficiently Blocks Rhinovirus Replication and Is Not Cross-Resistant with Rupintrivir. Antimicrob. Agents Chemother. 2015, 59, 5814–5818.

16. Lacroix, C.; George, S.; Leyssen, P.; Hilgenfeld, R.; Neyts, J. The enterovirus 3C protease inhibitor SG85 efficiently blocks rhinovirus replication and is not cross-resistant with rupintrivir. Antimicrob. Agents Chemother. 2015, 59, 5814–5818, doi:10.1128/AAC.00534-15.

17. Rocha-Pereira, J.; Nascimento, M.S.J.; Ma, Q.; Hilgenfeld, R.; Neyts, J.; Jochmans, D. The Enterovirus Protease Inhibitor Rupintrivir Exerts Cross-Genotypic Anti-Norovirus Activity and Clears Cells from the Norovirus Replicon. Antimicrob. Agents Chemother. 2014, 58, 4675–4681, doi:10.1128/AAC.02546-13.

18. Chmiela, S.; Sauceda, H.E.; Müller, K.-R.; Tkatchenko, A. Towards exact molecular dynamics simulations with machine-learned force fields. Nat. Commun. 2018, 9, 3887, doi:10.1038/s41467-018-06169-2.

19. Kawatkar, S.P.; Gagnon, M.; Hoesch, V.; Tiong-Yip, C.; Johnson, K.; Ek, M.; Nilsson, E.; Lister, T.; Olsson, L.; Patel, J.; et al. Design and structure-activity relationships of novel inhibitors of human rhinovirus 3C protease. Bioorg. Med. Chem. Lett. 2016, 26, 3248–3252, doi: http://dx.doi.org/10.1016/j.bmcl.2016.05.066.

20. Liu, Y.; Sheng, J.; Fokine, A.; Meng, G.; Shin, W.-H.; Long, F.; Kuhn, R.J.; Kihara, D.; Rossmann, M.G. Structure and inhibition of EV-D68, a virus that causes respiratory illness in children. Science 2015, 347, 71–74, doi:10.1126/science.1261962.

21. Pettersen, E.F.; Goddard, T.D.; Huang, C.C.; Couch, G.S.; Greenblatt, D.M.; Meng, E.C.; Ferrin, T.E. UCSF Chimera--a visualization system for exploratory research and analysis. J. Comput. Chem. 2004, 25, 1605–12, doi:10.1002/jcc.20084.

22. Hanwell, M.D.; Curtis, D.E.; Lonie, D.C.; Vandermeersch, T.; Zurek, E.; Hutchison, G.R. Avogadro: an advanced semantic chemical editor, visualization, and analysis platform. J. Cheminform. 2012, 4, 17, doi:10.1186/1758-2946-4-17.

23. Götz, A.W.; Williamson, M.J.; Xu, D.; Poole, D.; Le Grand, S.; Walker, R.C. Routine Microsecond Molecular Dynamics Simulations with AMBER on GPUs. 1. Generalized Born. J. Chem. Theory Comput. 2012, 8, 1542–1555, doi:10.1021/ct200909j.

24. Salomon-Ferrer, R.; Götz, A.W.; Poole, D.; Le Grand, S.; Walker, R.C. Routine Microsecond Molecular Dynamics Simulations with AMBER on GPUs. 2. Explicit Solvent Particle Mesh Ewald. J. Chem. Theory Comput. 2013, 9, 3878–3888, doi:10.1021/ct400314y.

25. Sprenger, K.G.; Jaeger, V.W.; Pfaendtner, J. The General AMBER Force Field (GAFF) Can Accurately Predict Thermodynamic and Transport Properties of Many Ionic Liquids. J. Phys. Chem. B 2015, 119, 5882–5895, doi:10.1021/acs.jpcb.5b00689.

26. Maier, J.A.; Martinez, C.; Kasavajhala, K.; Wickstrom, L.; Hauser, K.E.; Simmerling, C. ff14SB: Improving the Accuracy of Protein Side Chain and Backbone Parameters from ff99SB. J. Chem. Theory Comput. 2015, 11, 3696–3713, doi:10.1021/acs.jctc.5b00255.

27. Jorgensen, W.L.; Chandrasekhar, J.; Madura, J.D.; Impey, R.W.; Klein, M.L. Comparison of simple potential functions for simulating liquid water. J. Chem. Phys. 1983, 79, 926–935, doi:10.1063/1.445869.

28. Chetty, S.; Soliman, M.E.S. Possible allosteric binding site on Gyrase B, a key target for novel anti-TB drugs: homology modelling and binding site identification using molecular dynamics simulation and binding free energy calculations. Med. Chem. Res. 2015, 24, 2055–2074, doi:10.1007/s00044-014-1279-3.

29. Kumalo, H.M.; Soliman, M.E. A comparative molecular dynamics study on BACE1 and BACE2 flap flexibility. J. Recept. Signal Transduct. 2016, 36, 505–514, doi:10.3109/10799893.2015.1130058.

30. Roe, D.R.; Cheatham, T.E. PTRAJ and CPPTRAJ: Software for Processing and Analysis of Molecular Dynamics Trajectory Data. J. Chem. Theory Comput. 2013, 9, 3084–3095, doi:10.1021/ct400341p.

31. Kollman, P.A.; Massova, I.; Reyes, C.; Kuhn, B.; Huo, S.; Chong, L.; Lee, M.; Lee, T.; Duan, Y.; Wang, W.; et al. Calculating structures and free energies of complex molecules: combining molecular mechanics and continuum models. Acc. Chem. Res. 2000, 33, 889–897.

32. Massova, I.; Kollman, P.A. Combined molecular mechanical and continuum solvent approach (MM-PBSA/GBSA) to predict ligand binding. Perspect. Drug Discov. Des. 2000, 18, 113–135, doi:10.1023/A:1008763014207.

33. Kasahara, K.; Mohan, N.; Fukuda, I.; Nakamura, H. mDCC_tools: characterizing multi-modal atomic motions in molecular dynamics trajectories. Bioinformatics 2016, 32, 2531–2533, doi:10.1093/bioinformatics/btw129.

34. Desdouits, N.; Nilges, M.; Blondel, A. Principal Component Analysis reveals correlation of cavities evolution and functional motions in proteins. J. Mol. Graph. Model. 2015, 55, 13–24, doi:10.1016/j.jmgm.2014.10.011.

35. Martínez, L. Automatic Identification of Mobile and Rigid Substructures in Molecular Dynamics Simulations and Fractional Structural Fluctuation Analysis. PLoS One 2015, 10, e0119264, doi:10.1371/journal.pone.0119264.

36. Cocco, S.; Monasson, R.; Weigt, M. From Principal Component to Direct Coupling Analysis of Coevolution in Proteins: Low-Eigenvalue Modes are Needed for Structure Prediction. PLoS Comput Biol 2013, 9, e1003176.

37. Fuglebakk, E.; Echave, J.; Reuter, N. Measuring and comparing structural fluctuation patterns in large protein datasets. Bioinformatics 2012, 28, 2431–2440.

38. Blanco, F.; Ramírez-Alvarado, M.; Serrano, L. Formation and stability of beta-hairpin structures in polypeptides. Curr. Opin. Struct. Biol. 1998, 8, 107–111, doi:10.1016/S0959-440X(98)80017-1.

39. Lee, J.; Shin, S. Understanding β-Hairpin Formation by Molecular Dynamics Simulations of Unfolding. Biophys. J. 2001, 81, 2507–2516, doi:10.1016/S0006-3495(01)75896-1.

40. Diana, D.; De Rosa, L.; Palmieri, M.; Russomanno, A.; Russo, L.; La Rosa, C.; Milardi, D.; Colombo, G.; D’Andrea, L.D.; Fattorusso, R. Long range Trp-Trp interaction initiates the folding pathway of a pro-angiogenic β-hairpin peptide. Sci. Rep. 2015, 5, 16651, doi:10.1038/srep16651.

41. Gao, Y.; Li, Y.; Mou, L.; Lin, B.; Zhang, J.Z.H.; Mei, Y. Correct folding of an α-helix and a β-hairpin using a polarized 2D torsional potential. Sci. Rep. 2015, 5, 10359, doi:10.1038/srep10359.

42. Kaiser, L.; Crump, C.E.; Hayden, F.G. In vitro activity of pleconaril and AG7088 against selected serotypes and clinical isolates of human rhinoviruses. Antiviral Res. 2000, 47, 215–220, doi:http://dx.doi.org/10.1016/S0166-3542(00)00106-6.

43. Huang, Y.; Paul, D.R. Effect of molecular weight and temperature on physical aging of thin glassy poly(2,6-dimethyl-1,4-phenylene oxide) films. J. Polym. Sci. Part B Polym. Phys. 2007, 45, 1390–1398, doi:10.1002/polb.21173.

44. Pan, L.; Patterson, J.C. Molecular dynamics study of Zn(abeta) and Zn(abeta)2. PLoS One 2013, 8, e70681, doi:10.1371/journal.pone.0070681.

45. Kumalo, H.M.; Soliman, M.E. Per-Residue Energy Footprints-Based Pharmacophore Modeling as an Enhanced In Silico Approach in Drug Discovery: A Case Study on the Identification of Novel β-Secretase1 (BACE1) Inhibitors as Anti-Alzheimer Agents. Cell. Mol. Bioeng. 2016, 9, 175–189, doi:10.1007/s12195-015-0421-8.

46. Cele, F.N.; Ramesh, M.; Soliman, M.E.S. Per-residue energy decomposition pharmacophore model to enhance virtual screening in drug discovery: a study for identification of reverse transcriptase inhibitors as potential anti-HIV agents. Drug Des. Devel. Ther. 2016, 10, 1365–1377, doi:10.2147/DDDT.S95533.

